# Structural and Biophysical Basis for PFAS Binding by Human Sterol Carrier Protein-2

**DOI:** 10.1101/2025.10.27.684906

**Authors:** Aaron S. Birchfield, Rachel L. Signorelli, Kyla T. Cang, César A. Ramírez-Sarmiento, Brian Fuglestad

## Abstract

Per- and polyfluoroalkyl substances (PFAS) are harmful environmental contaminants that bioaccumulate in human tissues and are linked to adverse health outcomes. While PFAS are known to bind to a variety lipid binding proteins (LBPs), such as human serum albumin and fatty acid-binding proteins (FABPs), the broader molecular basis for their biological distribution and additional target proteins in humans remains unanswered. Motivated by its known promiscuity towards a range of hydrophobic ligands, we investigated the interaction between human sterol carrier protein 2 (SCP2) and various PFAS. SCP2 is a structurally distinct LBP with no previously reported affinity for PFAS. Using a combination of screening, fluorescence displacement assays, protein structure prediction of PFAS-SCP2 complexes, ITC, and NMR experiments, we demonstrate for the first time that SCP2 is a PFAS-binding protein. Our findings provide insight into the residues participating in these interactions and provide evidence for an additional LBP that may facilitate PFAS distribution and persistence in the human body.

---

Per- and polyfluoroalkyl substances (PFAS) are persistent synthetic chemicals that have been widely used for decades and are detected in water, soil, and air. They bioaccumulate in the liver, kidney, and testes, among other tissues, and persist in serum.^1–3^ Despite widespread exposure and links to negative health outcomes, little is known about cellular mechanisms of PFAS distribution. Evidence indicates that PFAS engage directly with proteins, raising critical questions about the molecular basis of their biological distribution, target proteins, and effects. To date, PFAS have been shown to bind a small yet diverse set of human proteins, including serum albumin, members of the fatty acid–binding protein (FABP) family, peroxisome proliferator-activated receptor γ, and transthyretin.^4–7^ A unifying feature among these proteins is the presence of large hydrophobic pockets that normally accommodate fatty acids or other lipids, offering plausible binding sites for PFAS. Reported PFAS binding affinities span from sub-micromolar to low millimolar, a range that is comparable to those of endogenous ligands.^4,5,8^ The emerging trend of protein types that bind to PFAS suggests that lipid-binding proteins (LBPs) may be particularly prone to bind PFAS. This study provides evidence of protein interactions that may be relevant to PFAS distribution and accumulation.

To provide further evidence for this model, we selected sterol carrier protein 2 (SCP2), a structurally distinct LBP with no prior PFAS interaction data. SCP2, also known as non-specific lipid-transfer protein, promiscuously binds and transports hydrophobic ligands, including cholesterol, long-chain fatty acids, acyl-CoA esters, and bile acids.^9–13^ The broad affinity to hydrophobic ligands, motivated us to investigate as a new PFAS-binding protein. Here we explore the link between PFAS chemical features and SCP2 binding and provide further evidence for the propensity of LBPs to engage with PFAS.

To assess the PFAS for SCP2 interactions, we performed a fluorescence displacement screen using fully delipidated, recombinant human SCP2 (Fig. S1) with NBD-stearic acid (NBD-SA) as a competitive probe. This fatty-acid analog is known to bind SCP2 with high affinity (K_d_ = 0.23-0.30 μM)^14,15^ and here exhibited a similar K_d_ of 0.31 μM (Fig. S2). PFAS were selected across major classes and screened at a fixed concentration of 10 μM (Fig. 1A-B). Fluorescence intensities were normalized, and compounds producing a reduction in signal to ≤0.75 (corresponding to ≥25% probe displacement) were designated as hits. Compounds meeting this criterion were advanced to competitive binding titrations to determine inhibition constants (K_i_).

**Fig. 1.**
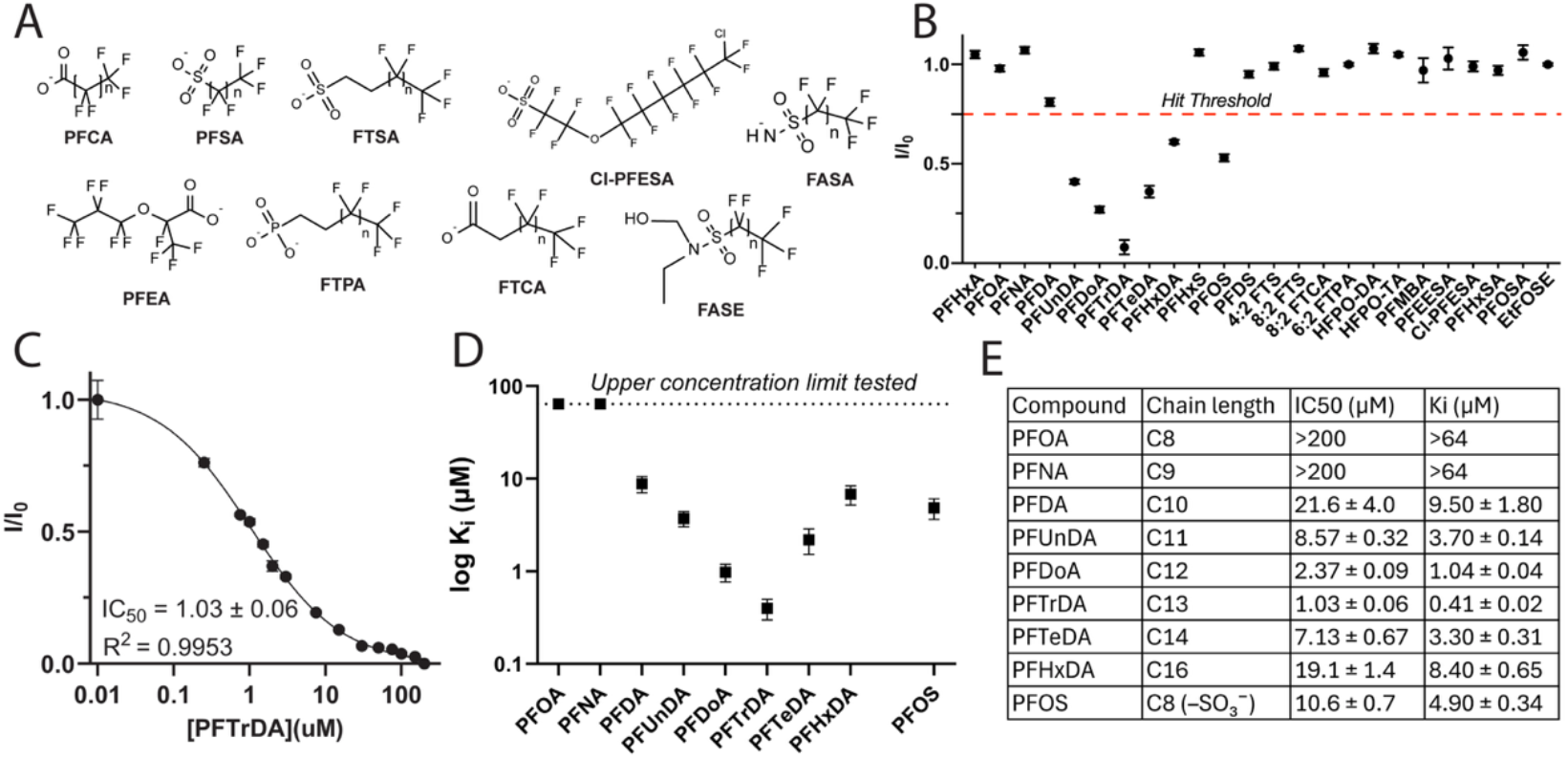
Tested PFAS, primary fluorescence displacement screen, and characterization of SCP2 binding to PFAS: (A) Chemical structures of PFAS classes included in the screening panel (full list and abbreviations in Table S1). (B) Competitive displacement screen of SCP2 using NBD-SA as a probe and a fixed 10 μM concentration of each PFAS. Data are shown as normalized fluorescence (I/I0). The red dashed line denotes the threshold used to designate screening hits advanced to affinity determination. (C) Fluorescence competition titration of PFTrDA displacing NBD-stearate from SCP2 (0.3 μM SCP2, 0.3 μM NBD-SA). (D) Affinity of PFAS hits from the SCP2 screen, plotted as log(K_i_). The dotted line marks the approximate upper detection limit of log(K_i_) under these assay conditions. (E) Summary of SCP2–PFAS potencies (IC_50_) and affinities (K_i_) from competition assays. Entries with no measurable displacement at the highest concentration are reported as lower bounds.

Fluorescent displacement assays of selected perfluoroalkyl carboxylic acids (PFCAs) revealed a chain-length dependence on affinity (Fig. 1C-D, Fig. S3). As chain length increased, affinity increased from PFDA (C10, K_i_ = 9.5 ± 1.8 μM) to a maximum at PFTrDA (C13, K_i_ = 0.41 ± 0.02 μM), then declined to PFHxDA (C16, K_i_ = 8.4 ± 0.6 μM) (Fig. 1D-E). This dependence suggests that longer chains enhance binding, beyond which additional carbons reduce affinity, plausibly due to steric constraints. Similar chain-length optima have been reported for FABP1 and FABP4.^4,16^Additionally, perfluorooctanesulfonic acid (PFOS) was shown to bind SCP2 with a K_i_ of 4.9 ± 0.3 μM (Fig. 1D-E, Fig. S4). Interestingly, the corresponding PFCA, PFOA, showed minimal binding up to 200 μM (Fig. 1D-E, Fig. S4), indicating that the sulfonate headgroup enables interactions not accessible to corresponding carboxylates. These results highlight two determinants of SCP2–PFAS recognition: chain length and headgroup chemistry. Reported SCP2 binding constants for lipid ligands span sub-micromolar to nanomolar values,^9,12,15,17,18^ with cholesterol binding with a K_d_ of 0.30 μM, similar to the tightest PFCA interactions observed here (e.g., PFTrDA, Ki = 0.41 μM).^17^ These comparisons indicate that the most strongly binding PFAS engage at affinities that overlap with endogenous ligands that are known to be transported by SCP2.

The fluorescence displacement assay provides a practical comparative screen for SCP2– PFAS interactions, but interpretation of IC_50_/K_i_ values carries important caveats. Variable Hill slopes across the PFAS series suggest non-ideal assay behavior rather than strict adherence to a simple one-site competitive binding model, potentially reflecting PFAS self-association, probe/ligand competition effects, or other deviations.^19–21^ In addition, for longer-chain PFAS, limited solubility and potential aggregation at higher concentrations may influence apparent displacement behavior. Dynamic light scattering was used to confirm a lack of major aggregation due to insolubility and revealed that the PFAS generally form micelles or other soluble assemblies under the assay conditions, as may be expected.^22,23^ SCP2 and other LBPs are known to interact with lipid assemblies to bind to hydrophobic ligands.^24^ Micellization may explain the non-ideal behavior of the displacement assays presented here and IC_50_/K_i_ should be interpreted comparatively rather than as a complete mechanistic description of binding.

To gain insight into binding mechanisms, we attempted to investigate PFCA binding to SCP2 by co-crystallography but were unsuccessful. This aligns with the longstanding challenges: to date, no ligand-bound mammalian SCP2 structures have been reported. Considering these limitations, we employed a deep-learning workflow (Boltz-2)^25^ for prediction of SCP2–PFCA complexes (C10–C16) (Fig. 2A). Confidence metrics indicate that the complex predictions are reliable, based on overall and protein-ligand interface predicted TM-scores (Table S2). Co-folding placed all PFCAs within the canonical hydrophobic pocket of SCP2,^26,27^ based on partial overlap with palmitate in holo-SCP2 from *Aedes aegypti* (PDB ID: 2KSI).^28^ In the predicted structures, the perfluorinated tail extends into the binding cavity and the carboxylate is oriented toward the solvent-exposed portal (Fig. 2A). This suggests that the portal region can stabilize polar headgroups, while the fluorinated tails are accommodated by hydrophobic β-sheet and helical cavity walls. This is consistent with mutagenesis in the portal that showed reduced long chain fatty acid (LFCA)/acyl-CoA binding,^29,30^ and suggests that PFCAs bind similarly at the LCFA binding site. These models provide valuable structural information and align with known fatty-acid/FA-CoA binding modes.

**Fig. 2.**
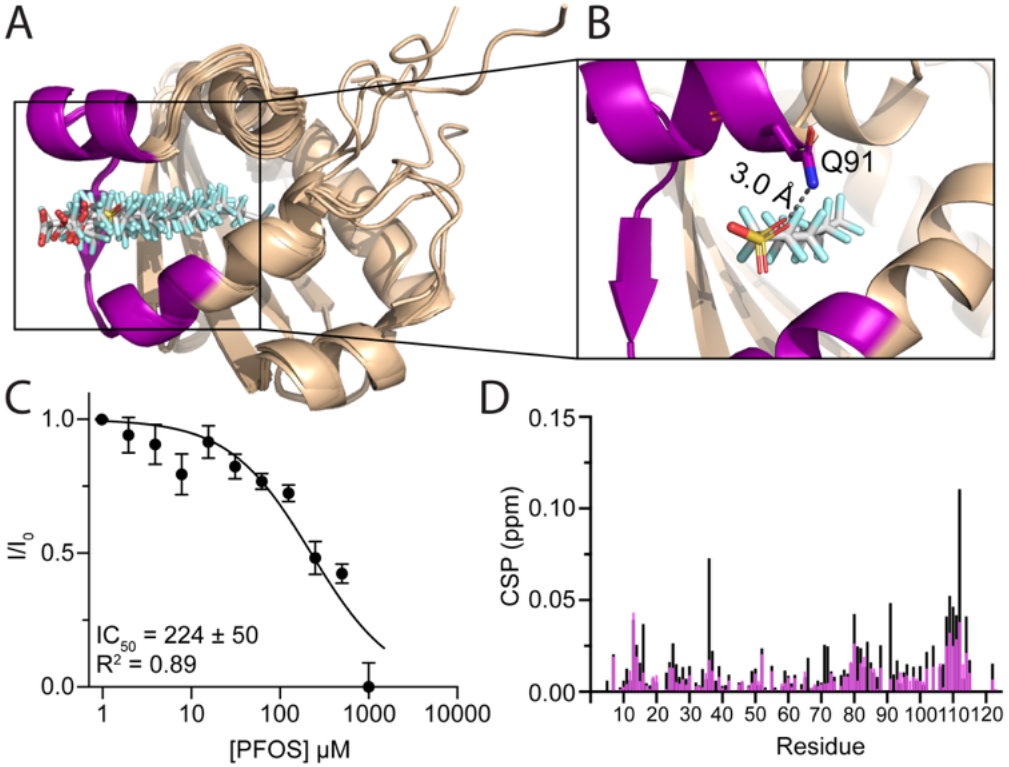
Deep learning-generated co-folding models of PFAS bound to SCP2 and mutational validation: (A) Boltz-2 co-folding models of SCP2 bound to long-chain PFCAs (C10-C16), with the portal region highlighted in purple. (B) Zoomed view of the portal region highlighting the proximity of Q91 to the sulfonate headgroup of PFOS. This residue was selected for mutation to glutamic acid to validate the model. (C) Binding of PFOS to the Q91E mutant is weaker than to wild-type SCP2 as measured by the NBD-SA displacement assay. IC_50_ is approximated as 224 ± 50 μM, which corresponds to a K_i_ of 82 ± 18 μM under these assay conditions. (D) CSPs of SCP2 WT with 1:1 molar ratio PFOS (black) compared to CSPs of SCP2 Q91E with 1:1 molar ratio PFOS (magenta). Corresponding NMR titrations in Fig. S6.

Using two different sets of parameters for the Boltz-2 co-folding predictions, increasing the recycling steps and diffusion samples from 3 and 5 (default) to 10 and 25 (exhaustive) led to similar binding poses (Fig. S5). The affinity determined by Boltz-2 did not correlate strongly with the experimental affinities from the displacement assays (Table S2). This is expected, as PFAS fall outside the chemical space represented in Boltz-2's training data, which is dominated by small molecules with hydrocarbon scaffolds. Per- and polyfluoroalkyl chains exhibit unique conformational and electrostatic properties, including the fluorine gauche effect, high rigidity, and atypical charge delocalization, which may not be well-represented in such a model.^31^ Regardless, confidence metrics for the predicted complex indicate reliable structural predictions (Table S2).

To validate the Boltz-2 modelling, we measured the effect of mutating a portal residue to weaken a SCP2-PFAS interaction. According to the models, the side-chain amide of Q91 is positioned near the PFAS headgroups (Fig. 2B). To induce repulsion with PFAS headgroups, we prepared a Q91E mutant and confirmed proper fold using protein NMR (Fig. S6). Binding of the NBD-SA probe was not weakened, consistent with the interactions of the long-tail fatty acid being primarily driven by hydrophobic interactions with the protein. Using the displacement assay, we found that the Q91E mutant binds to PFOS with a K_i_ = 82 ± 18 μM affinity, greater than 15-fold weaker than to wild-type SCP2 (Fig. 2C). This result was validated using protein NMR and chemical shift perturbation (CSP) mapping of PFOS binding to wild-type and Q91E SCP2, showing a muted binding effect upon mutation (Fig. 2D). This mutagenesis result confirms that the PFOS headgroup is positioned near Q91, helping to validate the Boltz-2 modeling results.

We next turned to protein NMR to verify SCP2–PFAS interactions and assess plausibility of the predictive models. CSP mapping of PFCAs binding to SCP2 was complicated by the required 5% ethanol co-solvent, whose presence partially mimicked the spectral changes of a liganded state. Mostly small PFCA-induced perturbations were observed (Figs. S7-S12), but CSP patterns varied somewhat across the PFCA series and should be interpreted cautiously in the presence of cosolvent. However, the overall perturbation pattern was consistent with the Boltz-2 models. The largest shifts occurred across the portal corridor, encompassing the β4–β5 region and adjacent a3–a5 helices, where the headgroup and tail contacts were predicted in the models. These data support SCP2 engagement by PFCAs but do not, on their own, unambiguously establish a single shared binding mode across the PFCA series.

To confirm modeling under fully aqueous conditions we used protein NMR to map CSPs upon PFOS binding (Fig. 3A), which needed no cosolvent. PFOS induced CSPs across the cavity and portal, spanning the β4–β5 and β1–β2 regions and adjacent a3–a5 helices (Fig. 3B). The largest shifts with CSPs greater than 0.1 include residues (E25, A36, D79, Q91, N104, A108, M109, L111, Q112, and L114). These CSPs correspond with residues previously identified at the cavity wall/portal by NMR^26,30^ and coincide with function-critical portal residues from mutagenesis studies.^29,30^ Altogether, the PFOS CSP data provide experimental support for PFOS engagement of the SCP2 cavity/portal region and support the plausibility of the PFOS predictive model.^10,17^ However, upon fitting the NMR titration data, a much weaker affinity was observed (global K_d_ = 227 ± 11 μM) compared to the displacement assay (Fig. 3C).

**Fig. 3.**
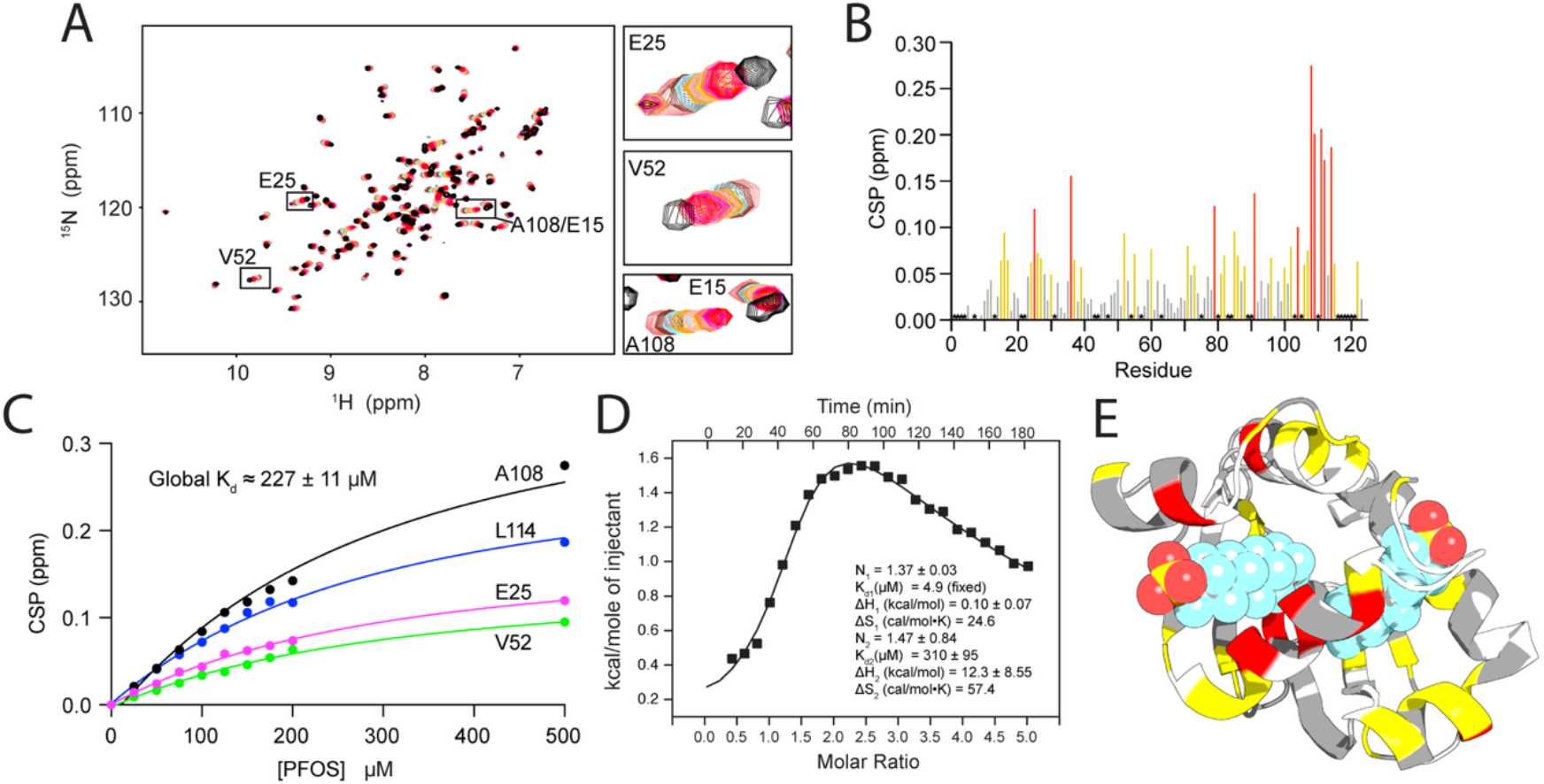
Structural analysis of SCP2–PFOS interactions. (A) ^1^H-^15^N HSQC titration of SCP2 apo (light coral) titrated with PFOS 0.25:1 (brown), 0.5:1 (cyan), 0.75:1 (orange), 1:1 (light pink), 1.25:1 (gold), 1.5:1 (coral), 1.75:1 (magenta), 2:1 (red), 5:1 (black) molar ratios of PFOS:SCP2. Zoomed panels are provided for some of the most shifted resonances. (B) CSPs of SCP2 upon PFOS addition, plotted versus residue number. Gold bars represent CSPs = 0.05 ppm, and red bars represent CSPs = 0.10 ppm. Gray bars represent CSPs < 0.05, while black stars represent residues that were overlapped or not observed. (C) CSPs of the highest shifting residues plotted against increasing PFOS concentrations to calculate the approximate K_d_ of the binding event of PFOS to SCP2 from NMR. (D) Two-site fitted ITC trace of PFOS (1 mM) being titrated into SCP2 (50 μM). (E) Boltz-2 model of SCP2–PFOS with CSP data from panel B mapped onto the structure (gray is < 0.05 ppm, yellow is = 0.05 ppm, red is = 0.10 ppm). White residues are unobserved due to lack of signal or overlap.

To reconcile this difference in affinities between assays, we performed isothermal titration calorimetry (ITC). We were able to obtain a binding isotherm for PFOS binding with SCP2 after subtraction of the background heat upon PFAS titration (Figs. 3D and S13). The longer chain PFCAs were attempted as well, however no interpretable ITC data were obtained, consistent with previous reports.^32^ The PFOS isotherm bore the signature of multiple, distinct binding events, which was best fit to a two-site binding model. To eliminate the uncertainty in the multi-parameter fits, the K_d_ of the higher affinity interaction (K_d1_) was fixed to the value obtained from the displacement assay (see supplement for details). A second, much weaker binding event was observed with a fitted K_d2_ = 310 ± 95 μM, correlating well with the NMR titration and suggesting that CSPs obtained via NMR are most sensitive to the second binding event. We note that the global fitting of the NMR data to obtain a K_d_ does not account for multiple binding events and should be considered an estimate. We also note that the second binding event has relatively large errors from the ITC fit from limitations on accessible PFOS concentrations points due to its solubility limits. Similar values were obtained while fixing the second binding site stoichiometry (N_2_) to that of the first (N_1_) (Table S3). Using this approach, a K_d1_ of 4.7 ± 0.8 μM was obtained, which correlates well with the displacement assay result. The ITC results provide strong validation for the K_d_ values obtained from the displacement assay for K_d1_, and the NMR titration result for K_d2_ for PFOS binding. These results show two distinct, entropically driven binding events (Fig. 3D). Modelling using Boltz-2 results in a plausible structural model for two PFOS binding sites that is consistent with NMR CSP data (Fig. 3E).

This work highlights SCP2 as a previously unknown PFAS-binding protein. A screen flagged long- and very-long-chain PFCAs and PFOS as hits; affinity characterization defined a chain-length dependence with a carbon chain length optimum of thirteen. The affinities determined here are within the range of affinities for other proteins binding to PFAS (Fig. 4, Table S4). For the PFAS characterized in this study, a range of nanomolar to millimolar affinities to target proteins have been reported, while we observed affinities in the low micromolar range to SCP2. This analysis indicates that SCP2 affinity for PFAS is significant compared to other known PFAS interacting proteins and suggests that SCP2 may have a role in biodistribution alongside other PFAS binding proteins. Boltz-2 models positioned PFAS within the lipid-binding SCP2 cavity, and the PFOS NMR spectra mapped perturbations to the portal and cavity-lining residues, mirroring the predicted pose. This work represents an application of the diffusion-based co-folding approach to PFAS–protein interactions and finds that, despite perfluoroalkyl chains lying outside the drug-like chemistries emphasized in current training sets, the approach can be used to rationalize ligand poses.

**Fig. 4.**
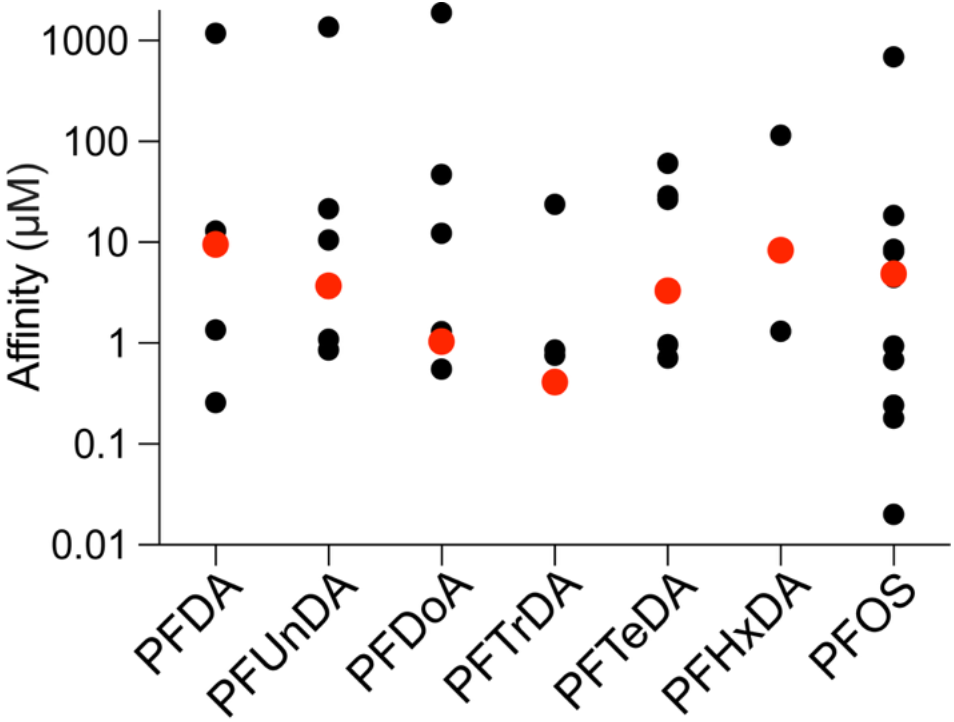
Comparison of SCP2-PFAS affinities to those of other reported PFAS binding proteins. Plot of K_i_ values determined in the current study via displacement assay (red) compared to published K_i_ or measured IC_50_ values of each PFAS binding to other proteins (black). Values, protein and PFAS identities, measurement methods, and citations for each data point are given in Table S4.

Discovery of SCP2 as a PFAS binding protein adds to a small but growing list of potential cellular carriers of these consequential toxicants. A small lipid-binding protein panel may provide useful for understanding PFAS-protein interactions. In the longer term, such a strategy could inform alternative fluoro-chemistries with reduced protein binding and potentially diminished biological residence and distribution. Integrating PFAS–protein interaction data with toxicological and exposure studies will be key to translating molecular binding into predictive insight on health risk and safer chemical design.

## Conflicts of interest

There are no conflicts to declare.

## Supporting Information

Additional experimental details and methods for SCP2 production, NBD–SA fluorescence displacement assays and analysis, NMR data for protein delipidation and PFAS binding, Boltz-2 protein–ligand modeling and ITC data; Tables S1–S4; Figures S1–S13; supplementary references (PDF). Boltz-2 predictive modelling data for this article are available at Zenodo at: https://doi.org/10.5281/zenodo.17438459.

## Author Contributions

A.S.B. and R.L.S. designed and performed experiments, analyzed and interpreted data, and wrote and edited the manuscript. K.T.C. performed experiments and edited the manuscript.

C.A.R.S. designed and performed calculations, analyzed and interpreted data, and wrote and edited the manuscript. B.F. designed experiments, interpreted data, and wrote and edited the manuscript.

## Acknowledgements

This work was funded by the National Institutes of Health (NIH) grant R35GM147221 and Agencia Nacional de Investigación y Desarrollo (ANID) through Fondo Nacional de Desarrollo Científico y Tecnológico (FONDECYT 1240205) and the ANID Millennium Science Initiative Program (ICN17_022). Powered@NLHPC: This research was partially supported by the supercomputing infrastructure of the NLHPC (CCSS210001). We gratefully acknowledge assistance from Dr. Yun Qu and Dr. Faik Musayev and helpful conversations from Dr. Elizabeth Komives and Dr. Kushol Gupta. CAR-S thanks the support from the Institute for Protein Design at the University of Washington on setting up Boltz-2 for the co-folding protein-ligand structure predictions.

## Notes

https://zenodo.org/records/17438460

### Competing Interest Statement

The authors have declared no competing interest.

### Summary of Updates

This revision includes new experiments, in particular ITC and NMR, that further investigates the interaction of PFOS with SCP2 and a potential second binding site. A mutagenesis experiment was performed that helped validate the structural model presented. An analysis of the affinity of the PFAS tested here versus other proteins, compared to their affinity to SCP2 is presented in a new figure. Edits to the text modify the discussion. Birchfield and Signorelli are designated as equal contributors in this version.

## References

(1) Maxwell, D. A. L., Oluwayiose, O. A., Houle, E., Roth, K., Nowak, K., Sawant, S., Paskavitz, A. L., Liu, W., Gurdziel, K., Petriello, M. C., and Richard Pilsner, J. (2024) Mixtures of per-and polyfluoroalkyl substances (PFAS) alter sperm methylation and long-term reprogramming of offspring liver and fat transcriptome. Environ. Int. 186, 108577.

(2) Vujic, E., Ferguson, S. S., and Brouwer, K. L. R. (2024) Effects of PFAS on human liver transporters: implications for health outcomes. Toxicological Sciences 200, 213.

(3) Wen, Y., Rashid, F., Fazal, Z., Singh, R., Spinella, M. J., and Irudayaraj, J. (2022) Nephrotoxicity of perfluorooctane sulfonate (PFOS)—effect on transcription and epigenetic factors. Environ. Epigenet. 8, dvac010.

(4) Birchfield, A. S., Musayev, F. N., Castillo, A. J., Zorn, G., and Fuglestad, B. (2025) Broad PFAS Binding with Fatty Acid Binding Protein 4 Is Enabled by Variable Binding Modes. JACS Au 5, 2469–2474.

(5) Jackson, T. W., Scheibly, C. M., Polera, M. E., and Belcher, S. M. (2021) Rapid Characterization of Human Serum Albumin Binding for Per- and Polyfluoroalkyl Substances Using Differential Scanning Fluorimetry. Environ. Sci. Technol.

(6) Pederick, J. L., Frkic, R. L., McDougal, D. P., and Bruning, J. B. (2024) A structural basis for the activation of peroxisome proliferator-activated receptor gamma (PPAR?) by perfluorooctanoic acid (PFOA). Chemosphere 354, 141723.

(7) Zhang, J., Begum, A., Brännström, K., Grundström, C., Iakovleva, I., Olofsson, A., Sauer-Eriksson, A. E., and Andersson, P. L. (2016) Structure-Based Virtual Screening Protocol for in Silico Identification of Potential Thyroid Disrupting Chemicals Targeting Transthyretin. Environ. Sci. Technol. 50, 11984–11993.

(8) Khazaee, M., Christie, E., Cheng, W., Michalsen, M., Field, J., and Ng, C. (2021) Perfluoroalkyl Acid Binding with Peroxisome Proliferator-Activated Receptors a, ?, and d, and Fatty Acid Binding Proteins by Equilibrium Dialysis with a Comparison of Methods. Toxics 2021, Vol. 9, Page 45 9, 45.

(9) Frolov, A., Cho, T. H., Billheimer, J. T., and Schroeder, F. (1996) Sterol carrier protein-2, a new fatty acyl coenzyme A-binding protein. Journal of Biological Chemistry 271, 31878–31884.

(10) Stolowich, N., Frolov, A., Petrescu, A. D., Scott, A. I., Billheimer, J. T., and Schroeder, F. (1999) Holo-sterol Carrier Protein-2: 13C NMR INVESTIGATION OF CHOLESTEROL AND FATTY ACID BINDING SITES. Journal of Biological Chemistry 274, 35425–35433.

(11) Li, N. C., Fan, J., and Papadopoulos, V. (2016) Sterol Carrier Protein-2, a Nonspecific Lipid-Transfer Protein, in Intracellular Cholesterol Trafficking in Testicular Leydig Cells. PLoS One 11, e0149728.

(12) Filipp, F. V., and Sattler, M. (2007) Conformational plasticity of the lipid transfer protein SCP2. Biochemistry 46, 7980–7991.

(13) Walters, S. H., Signorelli, R. L., Payne, A. G., Hojjatian, A., and Fuglestad, B. (2025) Compositional versatility enables biologically inspired reverse micelles for study of protein–membrane interactions. Soft Matter 21, 3547–3557.

(14) Schroeder, F., Myers-Payne, S. C., Billheimer, J. T., and Gibson Wood, W. (1995) Probing the ligand binding sites of fatty acid and sterol carrier proteins: effects of ethanol. Biochemistry 34, 11919–11927.

(15) Avdulov, N. A., Chochina, S. V., Igbavboa, U., Warden, C. S., Schroeder, F., and Wood, W. G. (1999) Lipid binding to sterol carrier protein-2 is inhibited by ethanol. Biochim. Biophys. Acta 1437, 37–45.

(16) Zhang, L., Ren, X. M., and Guo, L. H. (2013) Structure-based investigation on the interaction of perfluorinated compounds with human liver fatty acid binding protein. Environ. Sci. Technol. 47, 11293–11301.

(17) Colles, S. M., Woodford, J. K., Moncecchi, D., Myers-Payne, S. C., McLean, L. R., Billheimer, J. T., and Schroeder, F. (1995) Cholesterol interaction with recombinant human sterol carrier protein-2. Lipids 30, 795–803.

(18) Stolowich, N. J., Frolov, A., Atshaves, B., Murphy, E. J., Jolly, C. A., Billheimer, J. T., Ian Scott, A., and Schroeder, F. (1997) The sterol carrier protein-2 fatty acid binding site: an NMR, circular dichroic, and fluorescence spectroscopic determination. Biochemistry 36, 1719–1729.

(19) Prinz, H. (2009) Hill coefficients, dose–response curves and allosteric mechanisms. J. Chem. Biol. 3, 37.

(20) Hulme, E. C., and Trevethick, M. A. (2010) Ligand binding assays at equilibrium: validation and interpretation. Br. J. Pharmacol. 161, 1219–1237.

(21) Dong, D., Kancharla, S., Hooper, J., Tsianou, M., Bedrov, D., and Alexandridis, P. (2021) Controlling the self-assembly of perfluorinated surfactants in aqueous environments. Physical Chemistry Chemical Physics 23, 10029–10039.

(22) Kancharla, S., Jahan, R., Bedrov, D., Tsianou, M., and Alexandridis, P. (2021) Role of chain length and electrolyte on the micellization of anionic fluorinated surfactants in water. Colloids Surf. A Physicochem. Eng. Asp. 628, 127313.

(23) Allen, S. J., Dower, C. M., Liu, A. X., and Lumb, K. J. (2020) Detection of Small-Molecule Aggregation with High-Throughput Microplate Biophysical Methods. Curr. Protoc. Chem. Biol. 12, e78.

(24) Huang, H., Ball, J. M., Billheimer, J. T., and Schroeder, F. (1999) The sterol carrier protein-2 amino terminus: a membrane interaction domain. Biochemistry 38, 13231–13243.

(25) Passaro, S., Corso, G., Wohlwend, J., Reveiz, M., Thaler, S., Somnath, V. R., Getz, N., Portnoi, T., Roy, J., Stark, H., Kwabi-Addo, D., Beaini, D., Jaakkola, T., and Barzilay, R. (2025) Boltz-2: Towards Accurate and Efficient Binding Affinity Prediction. bioRxiv.

(26) García, F. L., Szyperski, T., Dyer, J. H., Choinowski, T., Seedorf, U., Hauser, H., and Wüthrich, K. (2000) NMR structure of the sterol carrier protein-2: implications for the biological role. J. Mol. Biol. 295, 595–603.

(27) Stanley, W. A., Filipp, F. V., Kursula, P., Schüller, N., Erdmann, R., Schliebs, W., Sattler, M., and Wilmanns, M. (2006) Recognition of a functional peroxisome type 1 target by the dynamic import receptor pex5p. Mol. Cell 24, 653–663.

(28) PDB-2ksi: Solution NMR structure of Sterol Carrier Protein - 2 from Aedes a… - Yorodumi.

(29) Seedorf, U., Scheek, S., Engel, T., Steif, C., Hinz, H. J., and Assmann, G. (1994) Structure-activity studies of human sterol carrier protein 2. Journal of Biological Chemistry 269, 2613–2618.

(30) Stolowich, N. J., Petrescu, A. D., Huang, H., Martin, G. G., Scott, A. I., and Schroeder, F. (2002) Sterol carrier protein-2: structure reveals function. Cell. Mol. Life Sci. 59, 193–212.

(31) Tiburtini, G. A., Bertarini, L., Bersani, M., Dragani, T. A., Rolando, B., Binello, A., Barge, A., and Spyrakis, F. (2024) In silico prediction of the interaction of legacy and novel per- and poly-fluoroalkyl substances (PFAS) with selected human transporters and of their possible accumulation in the human body. Arch. Toxicol. 98, 3035–3047.

(32) Crisalli, A. M., Cai, A., and Cho, B. P. (2023) Probing the interactions of perfluorocarboxylic acids of various chain lengths with human serum albumin: calorimetric and spectroscopic investigations. Chem. Res. Toxicol. 36, 703–713.

