## Supporting Information for "Structural and Biophysical Basis for PFAS Binding by Human Sterol Carrier Protein-2"

### Materials and Methods

#### Chemicals and Materials

Perfluorohexanoic Acid (PFHxA), Perfluorononanoic Acid (PFNA), Perfluorotetradecanoic Acid (PFTeDA), Perfluorotridecanoic Acid (PFTrDA), Perfluorododecanoic Acid (PFDoA), Perfluorodecanoic Acid (PFDA), Hexafluoropropylene Oxide Dimer Acid (HFPO-DA), Perfluorohexanesulfonamide (PFOSA), Perfluorohexanesulfonic acid (PFHxS), Perfluorodecanesulfonic acid (PFDS), 1H, 1H, 2H, 2H-Perfluorohexanesulfonic Acid (4:2 FTS) 1H, 1H, 2H, 2H-Perfluorodecane sulfonic acid (8:2 FTS), Hexafluoropropylene Oxide Dimer Acid (HFPO-DA), 9-chlorohexadecafluoro-3-oxanonane-1-sulfonic acid (6:2 Cl-PFESA), Perfluoro-4-methoxybutanoic Acid (PFMBA), Perfluorohexanesulfonamide (PFHxSA), Perfluorooctanesulfonamide (PFOSA), and N-Ethyl perfluorooctanesulfonamidoethanol (EtFOSE) were purchased from Cayman Chemical. Perfluorooctanesulfonic Acid (PFOS) and Perfluoro(2-ethoxyethane)sulfonic acid were purchased from Santa Cruz. Perfluorooctanoic Acid (PFOA) and 1H, 1H, 2H, 2H-Perfluorooctanephosphonic acid (6:2 FTPA) were obtained from Sigma-Aldrich. Perfluorohexadecanoic Acid (PFHxDA) and Hexafluoropropylene Oxide Trimer Acid (HFPO-TA) were acquired from Combi-Blocks. 12-NBD stearate, hereafter referred to as NBD-stearic acid (NBD-SA) was purchased from Avanti Polar Lipids. All other chemicals not explicitly mentioned in the text were purchased from commercial suppliers and used as received.

#### Protein Expression, Purification, and Delipidation

Human sterol carrier protein-2 (SCP2), wild-type and mutant Q91E, corresponding to UniProt P22307-2 (isoform 2) and to residues 425–547 of the canonical P22307 sequence, with the N-terminal mitochondrial transit peptide (residues 1–20) omitted, was expressed, purified, and delipidated as described in Walters *et al.* (2025).[1] Briefly, SCP2 containing an N-terminal poly-histidine affinity tag with a TEV protease cleavage site was expressed in BL21(DE3) *E. coli* using M9 minimal medium supplemented with $^{15}NH_4Cl$ for isotopic labeling. Protein expression was induced with 1 mM isopropyl-β-d-1-thiogalactopyranoside (IPTG) at 30 ºC overnight, and purification was performed by Ni-NTA affinity chromatography. The N-terminal tag was removed by TEV protease digestion, and the cleaved protein was further purified by a second Ni-NTA step. SCP2 has been shown to co-purify with endogenous *E. coli* lipids,[2] therefore, purified SCP2 was delipidated using Lipidex-5000 resin as described previously.[1] Delipidation was confirmed by $^{1}H$–$^{15}N$ HSQC NMR (Fig. S1) and the apo form was used for subsequent assays.

#### PFAS Fluorescence Displacement Assay

Fluorescence-based affinity assays for the NBD–SA probe were performed similarly to those previously described for ANS–FABP4 competition assays with modifications.[3] To measure probe affinity, 0.3 µM NBD–SA was incubated with increasing concentrations of SCP2 (0–3 µM) in 50 mM Tris–HCl (pH 7.5), 100 mM NaCl, 2 mM TCEP, and 0.5 mM EDTA at room temperature. After a 2 min equilibration in the dark, 250 µL aliquots were transferred to Greiner Black UV-STAR® 96 Well Microplates, and fluorescence was measured ($\lambda_{ex}$ = 467 nm, $\lambda_{em}$ = 538 nm) using a SpectraMax iD5 microplate reader. Data were fit by nonlinear regression using a quadratic ligand-depletion binding model in GraphPad Prism (Fig. S2).

Fluorescence displacement-based screening and competition assays were adapted from Birchfield *et al.* (2025) with modifications for SCP2 and NBD–SA as the probe.[3] Because most PFAS require a co-solvent for solubility,[3] assays were conducted in the presence of 5% ethanol or methanol, which produced only modest increases in the apparent $K_d$ of the NBD-SA probe (0.40 µM and 0.35 µM, respectively; Fig. S2). While SCP2 is known to weakly bind ethanol,[4,5] the minimal effect on NBD-SA affinity indicates minimal interference. To account for these effects, all controls and assays contained matched co-solvent, and $IC_{50}$ values were converted to $K_i$ using the probe $K_d$ measured under the solvent condition of the assay.

For the initial screening assay, concentrated PFAS stocks (200 µM) were prepared in either methanol or ethanol, depending on best solubility. A total of 15 µL of each stock was added to 275 µL of assay buffer (50 mM Tris-HCl pH 7.5, 100 mM NaCl, 2 mM TCEP, 0.5 mM EDTA) containing 0.3 µM SCP2 and 0.3 µM NBD–SA, yielding a final PFAS concentration of 10 µM and a uniform 5 % (v/v) co-solvent across all samples. For the 0 µM ligand control, 15 µL of co-solvent alone was added to account for solvent-related baseline effect. Samples were equilibrated for 2 min in the dark at room temperature, and 250 µL aliquots were transferred to Greiner Black UV-STAR® 96 Well Microplates for fluorescence measurement (λex = 467 nm, λem= 538 nm) on a SpectraMax iD5 plate reader. Each compound was tested in triplicate, and fluorescence intensities were normalized to the no-ligand control. Compounds producing ≥ 25 % displacement of NBD–SA from SCP2 in the presence of PFAS were designated as hits and advanced to follow-up titrations.

For affinity determination, SCP2 WT (0.3 µM) and NBD–SA (0.3 µM) were incubated with increasing concentrations of each PFAS hit using identical buffer and co-solvent conditions throughout at room temperature. SCP2 Q91E (0.3 µM) and NBD-SA (0.3 µM) were incubated with 1 mM PFOS and serial dilutions of PFOS were performed for affinity determination. All measured PFAS were tested with DLS at their highest concentrations under assay conditions, which confirmed that the samples were monodisperse and did not contain major aggregates. Fluorescence data were analyzed in GraphPad Prism using a sigmoidal four-parameter logistic (4PL) model (response vs. log[PFAS] to determine $IC_{50}$ values (Fig. S3 and S4), which were reported as $IC_{50}$ ± SE. Prism reports the standard error of log($IC_{50}$) because uncertainty is approximately symmetric in log-space. To express a single SE on the concentration scale, we applied the delta method, the standard first-order uncertainty propagation for $y = 10^x$ according to Taylor and Kuyatt (1994)[6]:

$$SE(IC_{50}) \approx ln\ ln\ 10\ * IC_{50} * SE(IC_{50})$$

$K_i$ values were computed from $IC_{50}$ using a free-concentration competitive model as implemented in a $IC_{50}$-to-$K_i$ converter.[7] The $K_d$ of NBD–SA used in these calculations corresponded to the solvent condition of each assay (0.31 µM with no co-solvent, 0.35 µM in 5 % MeOH, 0.40 µM in 5 % EtOH). Protein ($P$) and probe ($L$) concentrations were fixed at 0.3 µM as used experimentally. $K_i$ uncertainty was propagated from the $IC_{50}$ SE by first-order (delta-method) error propagation:

$$SE(K_i) \approx \frac{SE(IC_{50})}{D_{eff}}, \qquad D_{eff} = 1 + \frac{L_{50} + P_0}{K_d}$$

### NMR Spectroscopy

NMR samples contained $^{15}$N-labeled human SCP2 WT and Q91E were prepared as described above at 120 µM SCP2 in 20 mM Tris-HCl (pH 7.4), 100 mM NaCl, and 2 mM DTT, with 10 % (v/v) $D_2O$ as the lock solvent. Spectra were acquired at 37 °C on 600 or 700 MHz Bruker AVANCE III spectrometers equipped with a QXI probe. Assignments of the SCP2 protein were based on previously deposited assignments in the BMRB (Entry 4438)[8] and were confirmed and transferred using 3D HNCO, HNCACB, and CBCACONH NMR experiments on $^{15}$N-$^{13}$C labeled apo-SCP2 and 1.2:1 ligand bound SCP2.[9–11] PFAS (lyophilized or solvent-evaporated PFAS stocks) were added to the SCP2 sample to achieve a final 3-fold molar excess (360 µM PFAS). The following co-solvents were used at 5 % (v/v): methanol for PFUnDA, ethanol for PFDA, PFDoA, PFTrDA, and PFTeDA; no co-solvent was required for PFOS. Mixtures were sonicated for 1 h at room temperature prior to data collection to ensure equilibration. Two-dimensional $^{1}$H–$^{15}$N HSQC spectra were collected for each condition. Chemical shift perturbations (CSPs) were calculated as:

$$\Delta\delta = \sqrt{(\Delta^{1}H)^2 + \left(\frac{\Delta^{15}N}{9.8655}\right)^2}$$

NMR data were processed in NMRPipe and analyzed using NMRFAM-Sparky.[12,13]

### Protein-ligand predictions using Boltz-2

Protein-ligand complex structures were predicted using Boltz-2.[14] Boltz-2 is an end-to-end deep learning model that co-folds a protein sequence with a small molecule, represented by its SMILES string, to predict the three-dimensional structure of the bound complex. Additionally, Boltz-2 has been trained on binding affinity regression values and binary affinity classification data from different datasets to allow estimation of the binding energies of the ligands during the co-folding process.

For the co-folding predictions, the human SCP2 construct described above was input into Boltz-2 alongside the SMILES of different PFAS, including PFDA, PFDoA, PFHxDA, PFOS, PFTeDA, PFTrDA and PFUnDA, as well as palmitate to identify the binding pocket in comparison with the structure of *Aedes aegypti* SCP2 bound to palmitate (PDB ID: 2KSI).

For the protein-ligand complex predictions, two sets of parameters were employed for all cases. The first default set was performed with 3 recycling steps (--recycling_steps = 3) and 5 diffusion samples (--diffusion_samples = 5). The second exhaustive approach utilized 10 recycling steps (--recycling_steps = 10) and 25 diffusion samples (--diffusion_samples = 25). All predictions were executed using a Multiple Sequence Alignment (MSA) obtained from the MMseqs2 server with the --use_msa_server flag.[15]

The correlations between the estimated binding affinities from Boltz-2 using both sets of parameters and the ones obtained experimentally are shown in Supplementary Table S2. Given the similarities in the binding poses when using both sets of parameters (Supplementary Fig. S5), only predictions using 3 recycling steps and 10 diffusion samples are shown in Fig. 2.

All predicted protein-ligand complexes using Boltz-2 are available online in Zenodo (https://doi.org/10.5281/zenodo.17438460).

#### Isothermal Titration Calorimetry

ITC samples of SCP2 WT (50 μM) were dialyzed overnight before experiments in 20 mM Tris pH 7.4, 100 mM NaCl, 2 mM DTT and PFOS (1.36 mM) was prepared in the dialysate to avoid buffer mismatch. Titrations were performed on a MicroCal VP-ITC at 25 °C. Control titrations of buffer into PFOS were performed and the reference data was subtracted. Each titration consisted of 25 injections with the initial injection of 2 μL followed by 10 μL injections for the remaining titration points. Data was analyzed using Origin 7 with a MicroCal extension for ITC data processing. Each titration point was integrated, and the first injection point was removed before fitting the data. Data were fit to a one-site, two-sites, and sequential binding models in Origin, with the two-site model resulting in the best fit. Best fits were obtained with fixing $K_{d1}$ to the displacement assay values or fixing $N_2$ to the converged value of $N_1$, both with similar results. Allowing all parameters to float resulted in similar fit results but with very large errors.

**Table S1. PFAS compounds included in the SCP2 fluorescence displacement screen.** Compounds are organized by chemical class with common abbreviations and structures shown in the corresponding columns. Structures are depicted in the ionization states predicted to predominate at pH 7.4; all compounds are shown in their anionic forms except EtFOSE, which remains neutral under these conditions. Compound names follow conventional literature nomenclature (acid form).

| Chemical class | Compound name | Abbreviation | Structure |
|---|---|---|---|
| **Perfluoroalkyl Carboxylic Acids (PFCAs)** | Perfluorohexanoic acid | PFHxA | 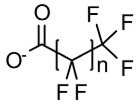 |
| | Perfluorooctanoic acid | PFOA | |
| | Perfluorononanoic acid | PFNA | |
| | Perfluorodecanoic acid | PFDA | |
| | Perfluoroundecanoic acid | PFUnDA | |
| | Perfluorododecanoic acid | PFDoA | |
| | Perfluorotridecanoic acid | PFTrDA | |
| | Perfluorotetradecanoic acid | PFTeDA | |
| | Perfluorohexadecanoic acid | PFHxDA | |
| **Perfluoroalkyl Sulfonic Acids (PFSAs)** | Perfluorohexanesulfonic acid | PFHxS | 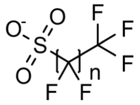 |
| | Perfluorooctanesulfonic acid | PFOS | |
| | Perfluorodecanesulfonic acid | PFDS | |
| **Fluorotelomer Sulfonic Acids** | 1H, 1H, 2H, 2H-Perfluorohexanesulfonic acid | 4:2 FTS | 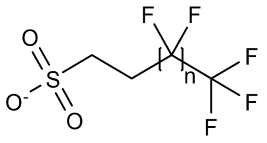 |
| | 1H, 1H, 2H, 2H-Perfluorodecanesulfonic acid | 8:2 FTS | |
| **Fluorotelomer Carboxylic Acid** | 2H, 2H-Perfluorodecanoic acid | 8:2 FTCA | 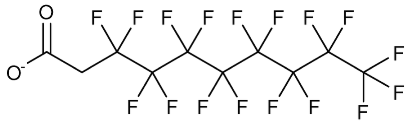 |
| **Fluorotelomer Phosphonic Acid** | 1H, 1H, 2H, 2H-Perfluorooctanephosphonic acid | 6:2 FTPA | 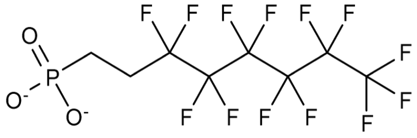 |
| **Perfluoroalkyl Ether Acids** | Hexafluoropropylene oxide dimer acid *(GenX)* | HFPO-DA | 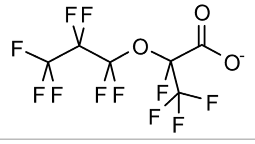 |

| Chemical class | Compound name | Abbreviation | Structure |
|---|---|---|---|
| | Hexafluoropropylene oxide trimer acid | HFPO-TA | 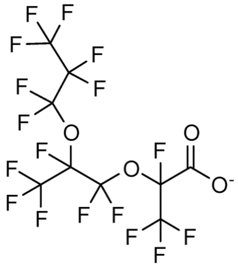 |
| | Perfluoro-4-methoxybutanoic acid | PFMBA | 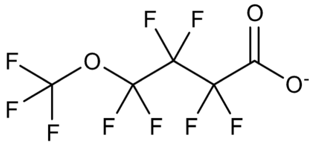 |
| | Perfluoro(2-ethoxyethane)sulfonic acid | PFEESA | 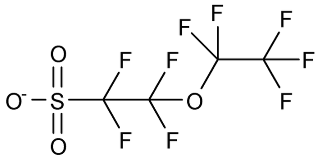 |
| **Chlorinated Perfluoroalkyl Ether Sulfonic Acid** | 9-Chlorohexadecafluoro-3-oxanonane-1-sulfonic acid *(F-53B)* | 6:2 Cl-PFESA | 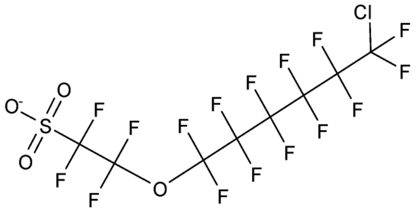 |
| **Perfluoroalkyl Sulfonamides** | Perfluorohexanesulfonamide | PFHxSA | 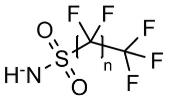 |
| | Perfluorooctanesulfonamide | PFOSA | |
| **Fluorooctane Sulfonamidoethanol** | N-Ethyl perfluorooctanesulfonamidoethanol | EtFOSE | 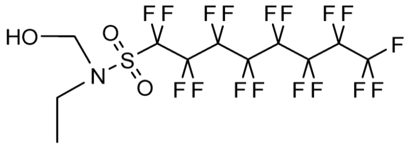 |

**Table S2**. **Predicted confidence metrics for Boltz-2 models of PFAS–SCP2 complexes generated under default (3 recycling / 5 diffusion) and exhaustive (10 recycling / 25 diffusion) sampling conditions.** Shown are overall and protein-ligand interface predicted TM-scores (pTM and ipTM), per-residue confidence metrics (complex pLDDT, complex iPLDDT), predicted distance errors (complex PDE, complex iPDE), and chain-specific pTM scores for each PFAS ligand (Chain A, SCP2; Chain B, ligand surrogate; Chain C, second ligand for SCP2 with 2 PFOS). Confidence Score refers to the overall Boltz-2 model confidence (0.8 * complex_plddt + 0.2 * iptm). Increasing recycling and diffusion depth produced minimal changes in global or interfacial confidence metrics.

| **PFAS** | **Recycling / Diffusion** | **pTM** | **ipTM** | **complex pLDDT** | **complex ipLDDT** | **complex PDE (Å)** | **complex iPDE (Å)** | **Chain A pTM** | **Chain B pTM (/Chain C pTM)** | **Confidence Score** |
|---|---|---|---|---|---|---|---|---|---|---|
| PFDA | 3 / 5 | 0.963 | 0.947 | 0.887 | 0.866 | 0.338 | 0.447 | 0.967 | 0.891 | 0.899 |
| PFDA | 10 / 25 | 0.940 | 0.916 | 0.893 | 0.872 | 0.428 | 0.645 | 0.947 | 0.868 | 0.898 |
| PFUnDA | 3 / 5 | 0.954 | 0.926 | 0.889 | 0.846 | 0.377 | 0.546 | 0.962 | 0.840 | 0.892 |
| PFUnDA | 10 / 25 | 0.963 | 0.947 | 0.891 | 0.881 | 0.340 | 0.454 | 0.968 | 0.862 | 0.902 |
| PFDoA | 3 / 5 | 0.954 | 0.931 | 0.882 | 0.855 | 0.369 | 0.534 | 0.962 | 0.849 | 0.892 |
| PFDoA | 10 / 25 | 0.943 | 0.916 | 0.886 | 0.861 | 0.413 | 0.623 | 0.953 | 0.836 | 0.892 |
| PFTrDA | 3 / 5 | 0.959 | 0.942 | 0.879 | 0.849 | 0.354 | 0.480 | 0.965 | 0.848 | 0.891 |
| PFTrDA | 10 / 25 | 0.950 | 0.929 | 0.881 | 0.847 | 0.378 | 0.536 | 0.958 | 0.844 | 0.891 |
| PFTeDA | 3 / 5 | 0.939 | 0.903 | 0.854 | 0.793 | 0.414 | 0.628 | 0.954 | 0.804 | 0.863 |
| PFTeDA | 10 / 25 | 0.946 | 0.914 | 0.866 | 0.813 | 0.392 | 0.580 | 0.959 | 0.827 | 0.875 |
| PFHxDA | 3 / 5 | 0.918 | 0.866 | 0.835 | 0.766 | 0.518 | 0.887 | 0.941 | 0.738 | 0.841 |
| PFHxDA | 10 / 25 | 0.945 | 0.919 | 0.862 | 0.811 | 0.407 | 0.615 | 0.957 | 0.805 | 0.874 |
| PFOS | 3 / 5 | 0.944 | 0.912 | 0.858 | 0.796 | 0.416 | 0.655 | 0.952 | 0.853 | 0.869 |
| PFOS | 10 / 25 | 0.925 | 0.894 | 0.857 | 0.782 | 0.463 | 0.727 | 0.934 | 0.857 | 0.865 |
| PFOSx2 | 3 / 5 | 0.946 | 0.913 | 0.787 | 0.661 | 0.472 | 0.769 | 0.962 | 0.876/0.875 | 0.812 |
| PFOSx2 | 10 / 25 | 0.957 | 0.934 | 0.797 | 0.688 | 0.421 | 0.648 | 0.968 | 0.897/0.888 | 0.824 |
| | | **min(pTM)** | **min(iPTM)** | **min(pLDDT)** | **min(ipLDDT)** | **max(PDE)** | **max(iPDE)** | **min(chainPTM)** | **min(chainPTM)** | **min(CS)** |
| | 3 / 5 | 0.918 | 0.866 | 0.787 | 0.661 | 0.518 | 0.887 | 0.941 | 0.738 | 0.812 |
| | 10 / 25 | 0.925 | 0.894 | 0.797 | 0.688 | 0.463 | 0.727 | 0.934 | 0.805 | 0.824 |

**Table S3. Results of two-site fitting of ITC data for PFOS-SCP2 titration while fixing the $N_2$ to match the converged $N_1$ value.** Resulting fitted curve is indistinguishable from that depicted in Fig. 3D.

| | |
|---|---|
| $N_1$ | 1.37 ± 0.03 |
| $K_{d1}$ (μM) | 4.7 ± 0.8 |
| $\Delta H_1$ (kcal/mol) | 0.12 ± 0.08 |
| $\Delta S_1$ (cal/mol·K) | 24.8 |
| $N_2$ | 1.37 (fixed) |
| $K_{d2}$ (μM) | 327 ± 44 |
| $\Delta H_2$ (kcal/mol) | 13.6 ± 1.0 |
| $\Delta S_2$ (cal/mol·K) | 57.4 |

**Table S4.** $K_i$ or $IC_{50}$ values for protein interactions with the PFAS characterized in this study. Protein abbreviations are: HSA, human serum albumin; FABP4, fatty acid binding protein 4; FABP1, fatty acid binding protein 1; TTR, transthyretin; PPARγ, peroxisome proliferator-activated receptor γ; PPARδ, peroxisome proliferator-activated receptor δ; TBG, thyroxine-binding globulin.

| PFAS | Reported Ki or IC50 | Reference | Protein | Method |
|---|---|---|---|---|
| PFDA | 1190 | 16 | HSA | Differential Scanning Fluorimetry |
| | 1.35 | 3 | FABP4 | Fluorescence competition |
| | 12.9 | 17 | FABP1 | Fluorescence competition |
| | 0.259 | 18 | TTR | Fluorescence competition |
| | 8.954 | 19 | TTR | Radioligand competition |
| PFDoA | 1890 | 16 | HSA | Differential Scanning Fluorimetry |
| | 0.551 | 3 | FABP4 | Fluorescence competition |
| | 12.3 | 17 | FABP1 | Fluorescence competition |
| | 1.293 | 18 | TTR | Fluorescence competition |
| | 46.894 | 19 | TTR | Radioligand competition |
| PFHxDA | 1.312 | 3 | FABP4 | Fluorescence competition |
| | 115.4 | 17 | FABP1 | Fluorescence competition |
| PFOS | 690 | 16 | HSA | Differential Scanning Fluorimetry |
| | 0.18 | 20 | TTR | Fluorescence competition |
| | 8.5 | 20 | PPARγ | Equilibrium dialysis |
| | 0.686 | 20 | PPARδ | Equilibrium dialysis |
| | 4.504 | 3 | FABP4 | Fluorescence competition |
| | 0.243 | 21 | TTR | Fluorescence competition |
| | 18.5 | 17 | FABP1 | Fluorescence competition |
| | 0.02 | 18 | TTR | Fluorescence competition |
| | 0.94 | 19 | TTR | Radioligand competition |
| | 8.1 | 22 | FABP1 | Fluorescence competition |
| PFTeDA | 0.7126 | 3 | FABP4 | Fluorescence competition |
| | 60.5 | 17 | FABP1 | Fluorescence competition |
| | 26.6 | 18 | TBG | Fluorescence competition |
| | 0.972 | 18 | TTR | Fluorescence competition |
| | 28.996 | 19 | TTR | Radioligand competition |
| PFTrDA | 0.757 | 3 | FABP4 | Fluorescence competition |
| | 23.8 | 18 | TBG | Fluorescence competition |
| | 0.858 | 18 | TTR | Fluorescence competition |
| PFUnDA | 1.094 | 3 | FABP4 | Fluorescence competition |
| | 10.6 | 17 | FABP1 | Fluorescence competition |
| | 0.854 | 18 | TTR | Fluorescence competition |
| | 21.56 | 19 | TTR | Radioligand competition |
| | 1360 | 16 | HSA | Differential Scanning Fluorimetry |

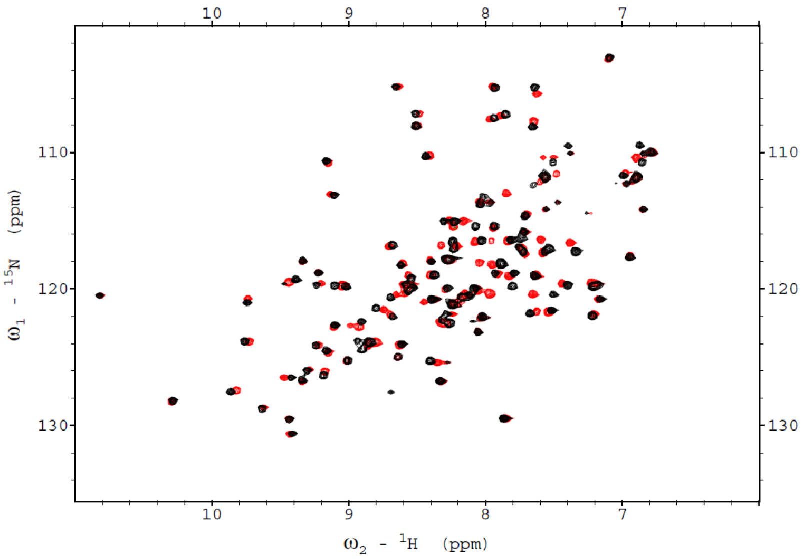

**Fig. S1. Verification of SCP2 delipidation by $^{1}H$–$^{15}N$ HSQC NMR.** Overlay of $^{1}H$–$^{15}N$ HSQC spectra of delipidated (apo) SCP2 (red) and relipidated SCP2 bound to palmitic acid (black) collected at pH 7.4 and 37 °C. The spectral differences between the apo and holo forms confirm successful delipidation of recombinant SCP2. Spectra were acquired on a 600 MHz Bruker AVANCE III spectrometer in 20 mM Tris-HCl, 100 mM NaCl, and 2 mM DTT, processed in NMRPipe, and analyzed using NMRFAM-Sparky.[12,13]

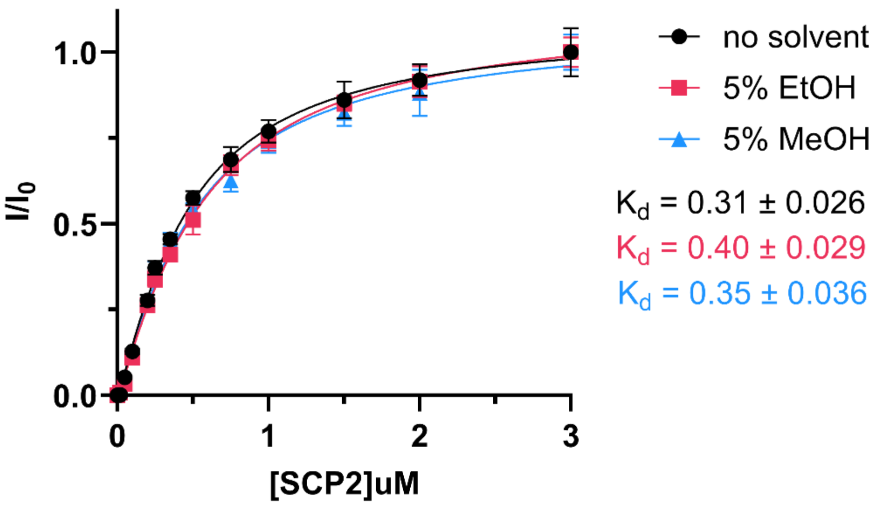

**Fig. S2. Determination of the apparent $K_d$ of NBD-SA for SCP2 under varying solvent conditions.** Binding of NBD-stearic acid with SCP2 was measured under three solvent conditions: no solvent ($K_d$ = 0.31 μM), 5% ethanol ($K_d$ = 0.40 μM), and 5% methanol ($K_d$ = 0.35 μM). Data were fit using a quadratic ligand depletion binding model. Solvent inclusion caused only minor increases in apparent $K_d$, which were accounted for when converting $IC_{50}$ to $K_i$ in displacement assays. Data points represent triplicate measurements, with error bars indicating the standard deviation.

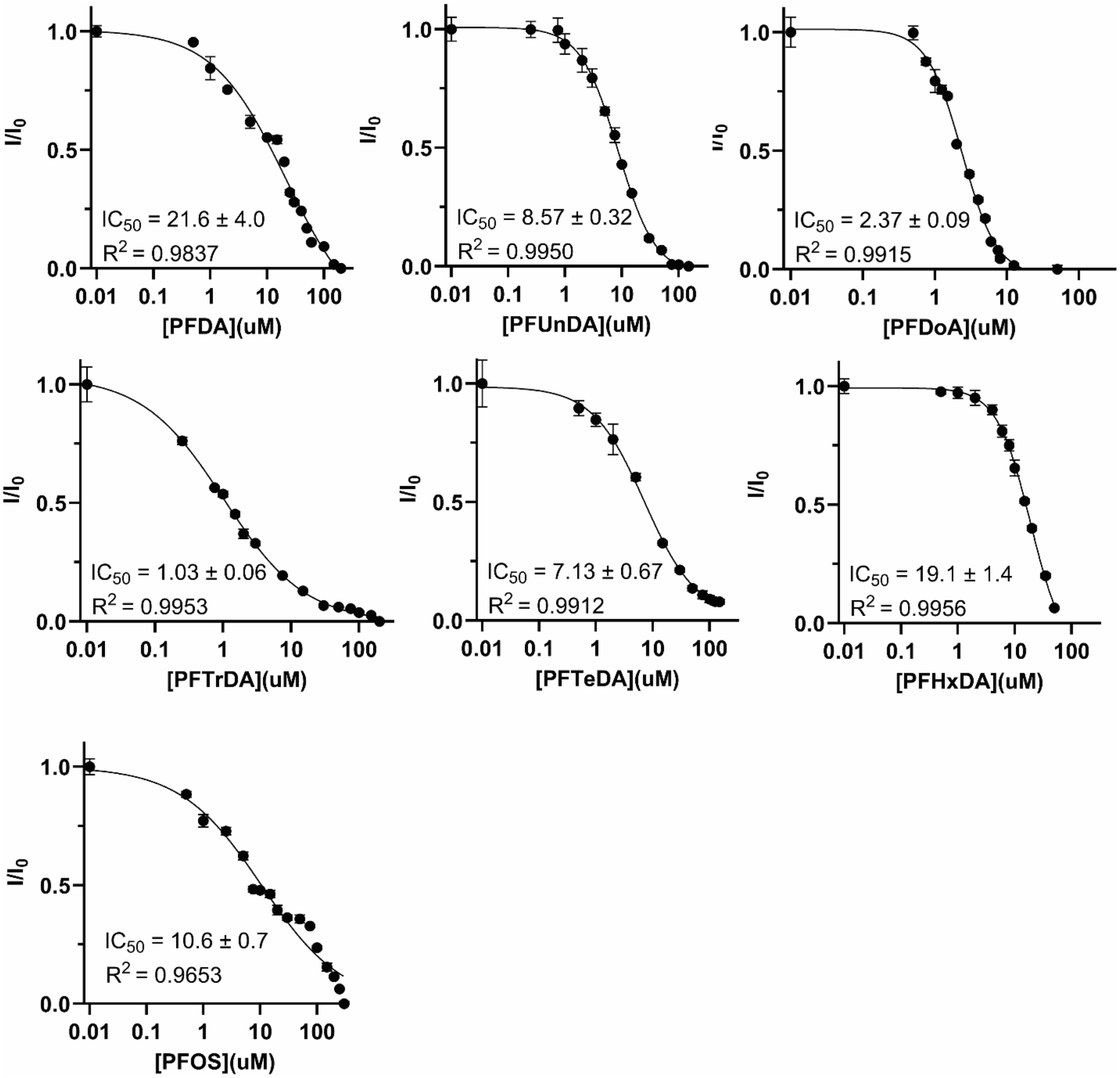

**Fig. S3. SCP2–PFAS fluorescence competition assays**. Representative fluorescence competition titrations of SCP2 (0.3 μM) with NBD-stearic acid (0.3 μM) in the presence of increasing concentrations of PFAS. $IC_{50}$ values (± SEM) were determined by nonlinear regression using a four-parameter model in GraphPad Prism. Data points represent triplicate measurements, with error bars indicating the standard deviation.

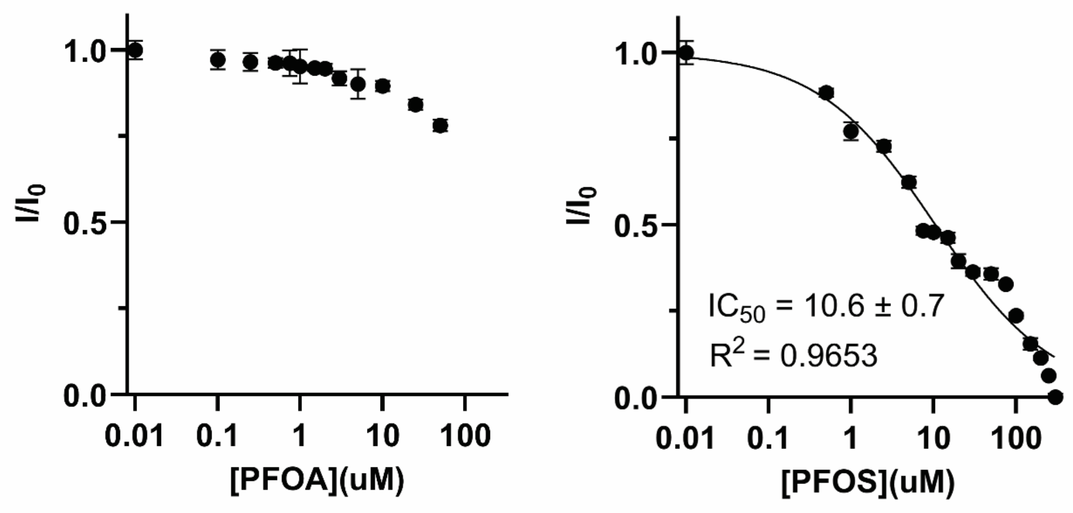

**Fig. S4. Comparison of SCP2 inhibition by PFOA and PFOS.** Fluorescence competition curves for SCP2 with perfluorooctanoic acid (PFOA, left) and perfluorooctanesulfonic acid (PFOS, right), obtained using 0.3 µM SCP2 and 0.3 µM NBD-stearic acid in 50 mM Tris-HCl pH 7.4, 100 mM NaCl, and 2 mM TCEP at 25 °C. PFOS produced a clear, saturable dose–response with an $IC_{50}$ = 10.6 ± 0.7 µM ($R^2$ = 0.965), whereas PFOA showed markedly less binding under similar conditions.

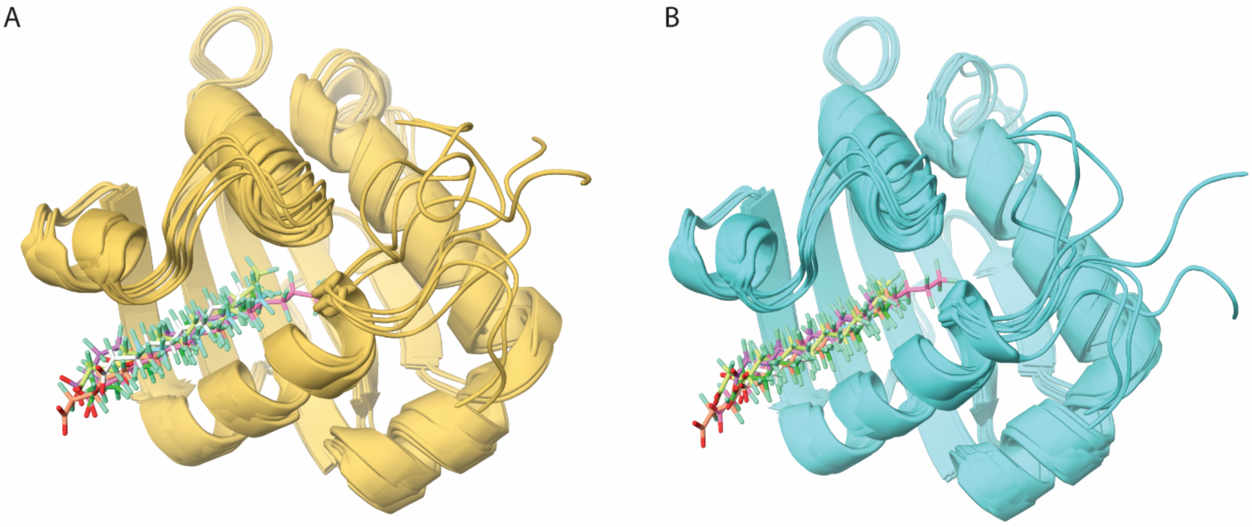

**Fig. S5. Effect of increased recycling and diffusion sampling on Boltz-2 predictions of PFAS-bound SCP2.** (A) Ensemble of Boltz-2 models for PFCA C10–C16 and PFOS generated with a default protocol of 3 recycling steps and 5 diffusion samples. (B) Ensemble of corresponding models generated with an exhaustive protocol of 10 recycling steps and 25 diffusion samples. Increasing the number of recycling iterations and diffusion samples produced highly similar binding poses, indicating convergence of the predicted PFAS-binding mode.

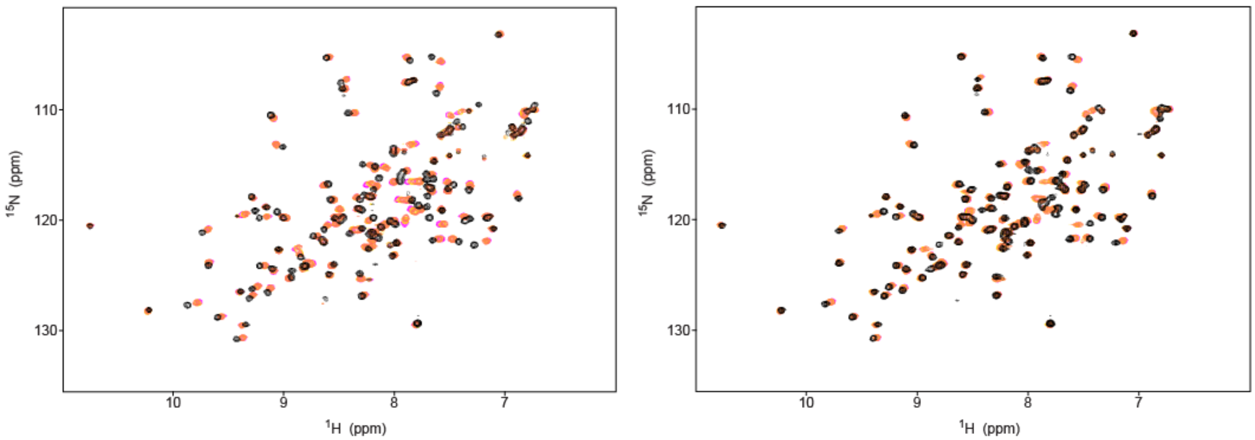

**Fig. S6. NMR titrations comparing SCP2 WT vs. SCP2 Q91E with PFOS.** SCP2 WT NMR titration with PFOS (left) and SCP2 Q91E NMR titration with PFOS (right). Left: $^{1}H$–$^{15}N$ HSQC spectra of apo SCP2 WT (magenta), 0.5:1 molar ratio PFOS:SCP2 WT (gold), 1:1 molar ratio PFOS:SCP2 WT(coral), 5:1 molar ratio PFOS:SCP2 WT(black). Right: $^{1}H$–$^{15}N$ HSQC spectra of apo SCP2 Q91E (magenta), 0.5:1 molar ratio PFOS:SCP2 Q91E (gold), 1:1 molar ratio PFOS:SCP2 Q91E (coral), 5:1 molar ratio PFOS:SCP2 Q91E (black).

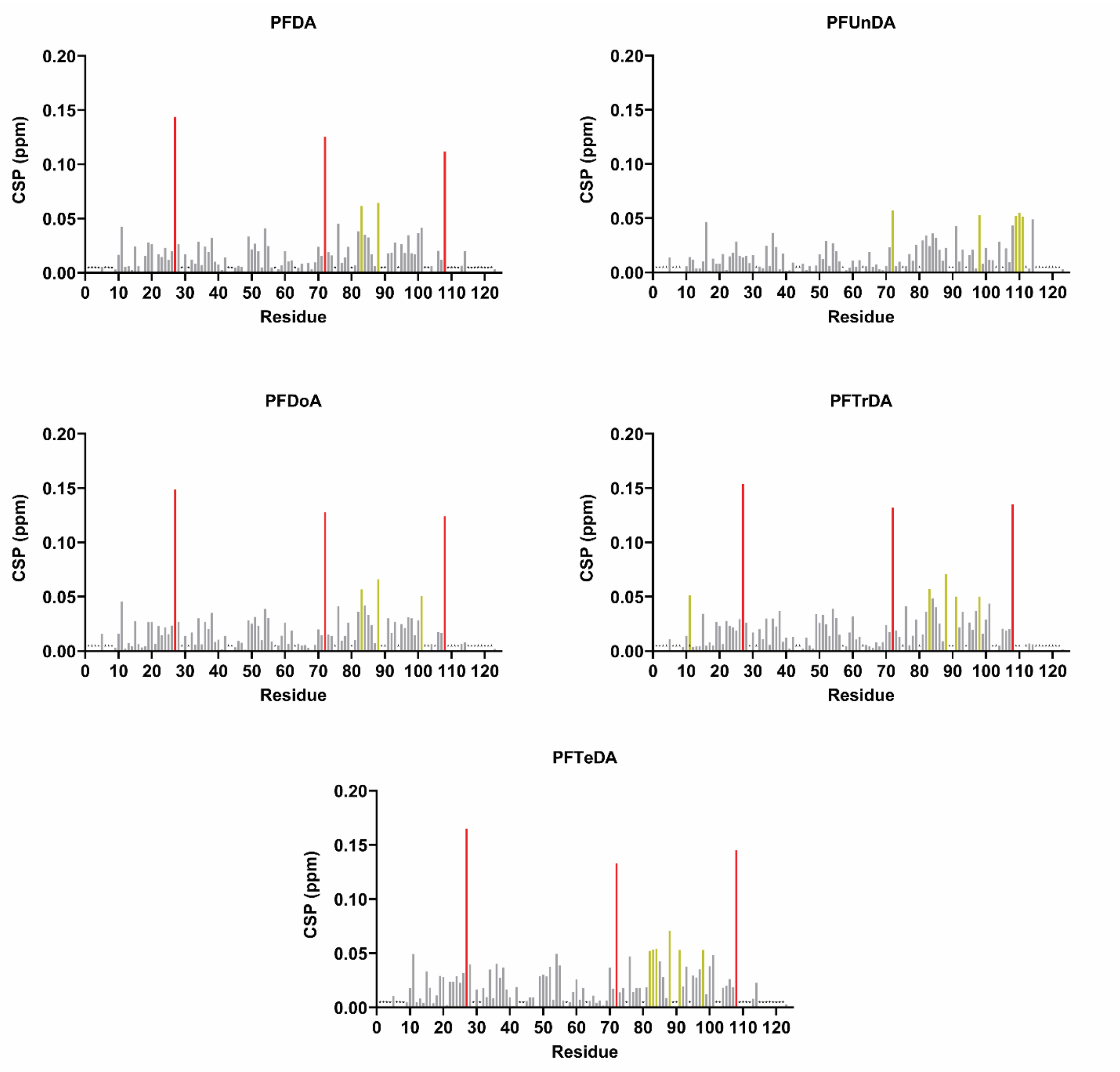

**Fig. S7. Chemical shift perturbations of SCP2 induced by PFCA binding.** Residue-specific $^{1}$H–$^{15}$N HSQC chemical shift perturbations (CSPs) for SCP2 titrated with perfluorinated carboxylic acids (PFCAs) of increasing chain length—PFDA, PFUnDA, PFDoA, PFTrDA, and PFTeDA—are shown. Gold bars represent residues with CSPs ≥ 0.05 ppm and < 0.10 ppm, red bars represent CSPs ≥ 0.10 ppm, and gray bars represent CSPs < 0.05 ppm. Black stars denote residues for which CSPs could not be derived. These data highlight perturbation patterns across the PFCA series, with strongest effects localized near the canonical fatty-acid binding site.

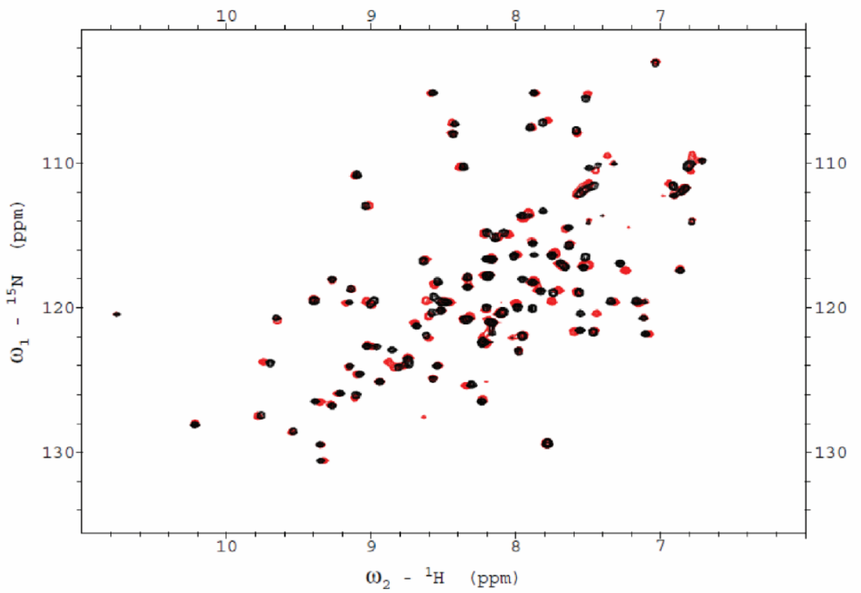

**Fig. S8. Overlay of $^{1}$H–$^{15}$N HSQC spectra of SCP2 with Perfluorodecanoic Acid (PFDA).** $^{1}$H–$^{15}$N HSQC spectra of apo SCP2 (red) and SCP2 in complex with PFDA (black) at a 1:3 SCP2:PFDA molar ratio. Shifts in cross-peaks indicate residue-specific perturbations consistent with ligand binding within the hydrophobic cavity.

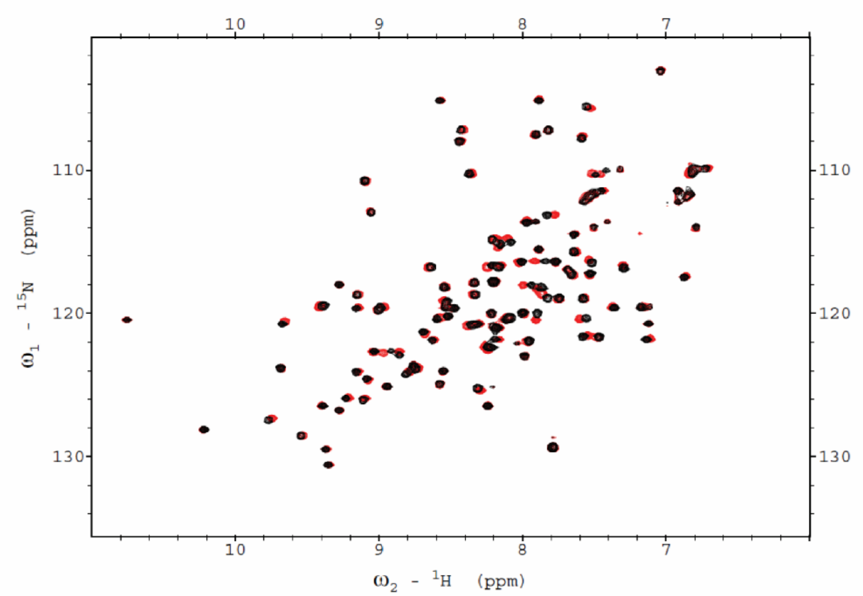

**Fig S9. Overlay of $^{1}$H–$^{15}$N HSQC spectra of SCP2 with Perfluoroundecanoic Acid (PFUnDA).** $^{1}$H–$^{15}$N HSQC spectra of apo SCP2 (red) and SCP2 in complex with PFUnDA (black) at a 1:3 SCP2:PFUnDA molar ratio. Shifts in cross-peaks indicate residue-specific perturbations consistent with ligand binding within the hydrophobic cavity.

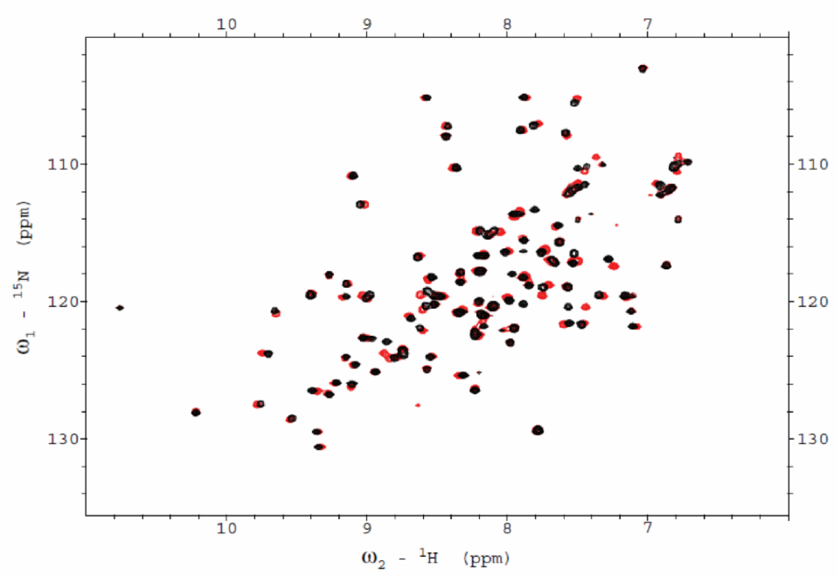

**Fig. S10. Overlay of $^{1}H$–$^{15}N$ HSQC spectra of SCP2 with Perfluorododecanoic Acid (PFDoA).** $^{1}H$–$^{15}N$ HSQC spectra of apo SCP2 (red) and SCP2 in complex with PFDoA (black) at a 1:3 SCP2:PFDoA molar ratio. Shifts in cross-peaks indicate residue-specific perturbations consistent with ligand binding within the hydrophobic cavity.

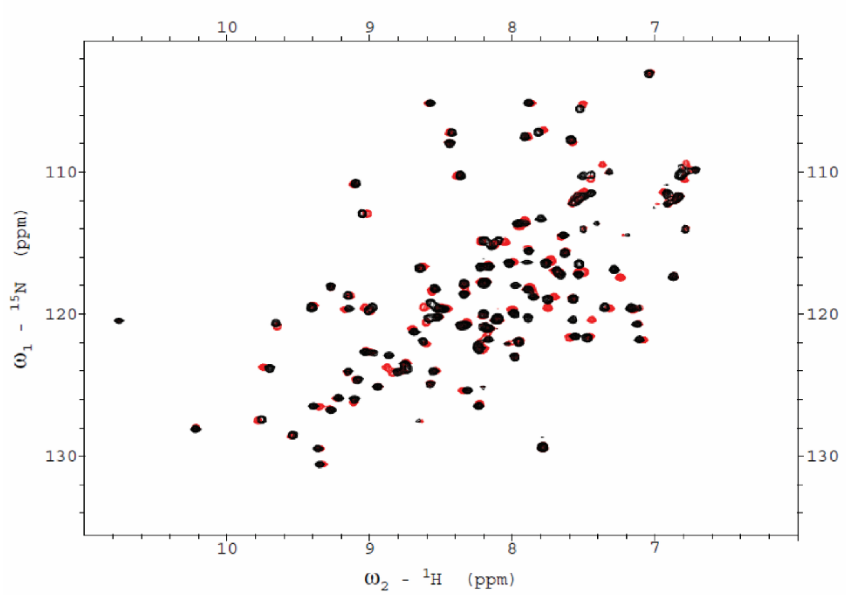

**Fig. S11. Overlay of $^1$H–$^{15}$N HSQC spectra of SCP2 with Perfluorotridecanoic Acid (PFTrDA).** $^1$H–$^{15}$N HSQC spectra of apo SCP2 (red) and SCP2 in complex with PFTrDA (black) at a 1:3 SCP2:PFTrDA molar ratio. Shifts in cross-peaks indicate residue-specific perturbations consistent with ligand binding within the hydrophobic cavity.

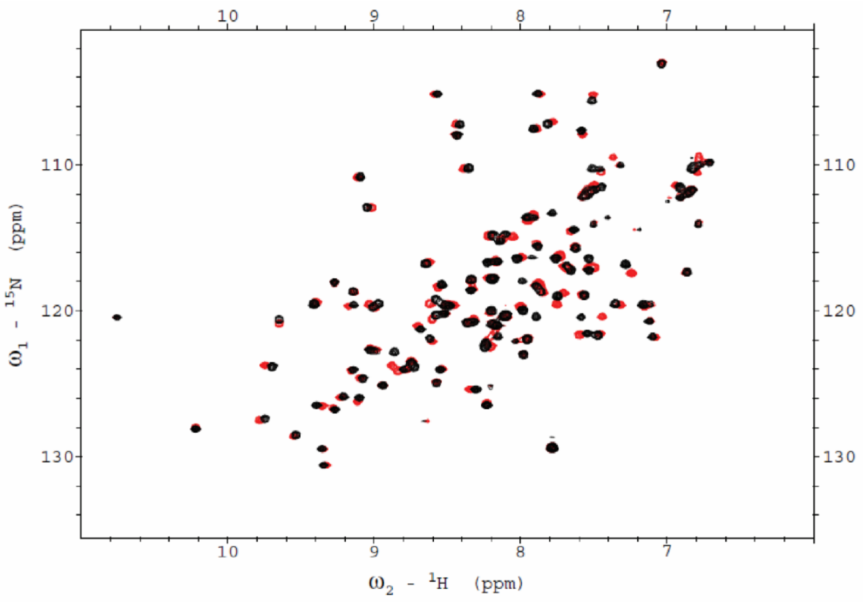

**Fig. S12. Overlay of $^{1}$H–$^{15}$N HSQC spectra of SCP2 with Perfluorotetradecanoic Acid (PFTeDA).** $^{1}$H–$^{15}$N HSQC spectra of apo SCP2 (red) and SCP2 in complex with PFTeDA (black) at a 1:3 SCP2:PFTeDA molar ratio. Shifts in cross-peaks indicate residue-specific perturbations consistent with ligand binding within the hydrophobic cavity.

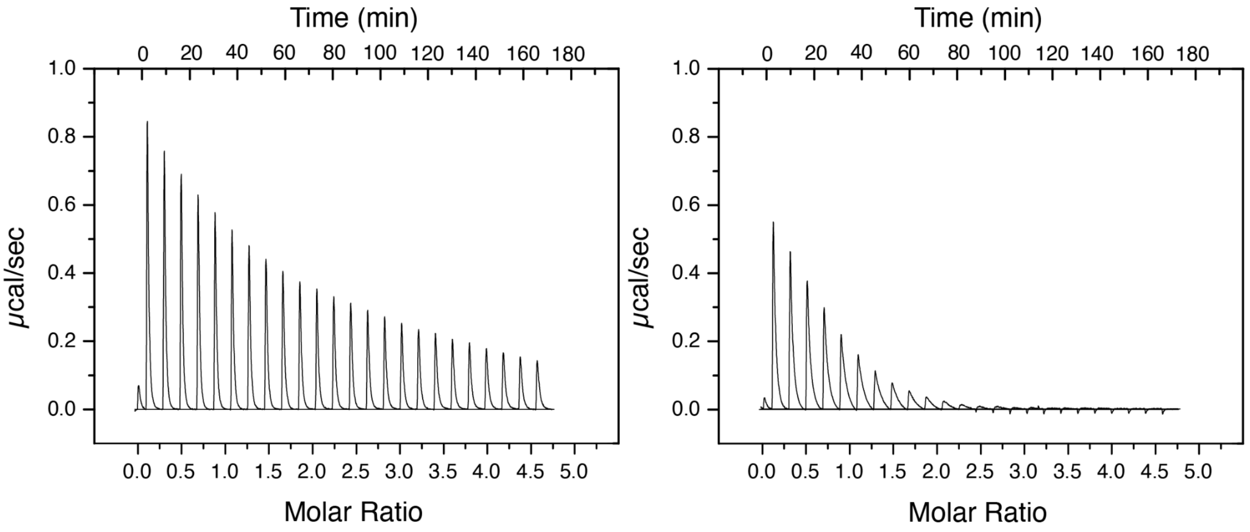

**Fig S13. Isothermal titration calorimetry with SCP2 WT and PFOS.** ITC thermograms showing the raw traces of 1.36 mM PFOS being titrated into 50 μM SCP2 (left) and the control of 1.36 mM PFOS being titrated into buffer (right).

**Supplemental References**

(1) Walters, S. H.; Signorelli, R. L.; Payne, A. G.; Hojjatian, A.; Fuglestad, B. Compositional Versatility Enables Biologically Inspired Reverse Micelles for Study of Protein–Membrane Interactions. *Soft Matter* **2025**, *21*, 3547–3557.

(2) Stolowich, N. J.; Frolov, A.; Atshaves, B.; Murphy, E. J.; Jolly, C. A.; Billheimer, J. T.; Scott, A. I.; Schroeder, F. The Sterol Carrier Protein-2 Fatty Acid Binding Site: An NMR, Circular Dichroic, and Fluorescence Spectroscopic Determination. *Biochemistry* **1997**, *36* (7), 1719–1729.

(3) Birchfield, A. S.; Musayev, F. N.; Castillo, A. J.; Zorn, G.; Fuglestad, B. Broad PFAS Binding with Fatty Acid Binding Protein 4 Is Enabled by Variable Binding Modes. *JACS Au* **2025**, *5* (6), 2469–2474. https://doi.org/10.1021/jacsau.5c00504.

(4) Avdulov, N. A.; Chochina, S. V; Igbavboa, U.; Warden, C. S.; Schroeder, F.; Wood, W. G. Lipid Binding to Sterol Carrier Protein-2 Is Inhibited by Ethanol. *Biochimica et Biophysica Acta (BBA)-Molecular and Cell Biology of Lipids* **1999**, *1437* (1), 37–45.

(5) Schroeder, F.; Myers-Payne, S. C.; Billheimer, J. T.; Wood, W. G. Probing the Ligand Binding Sites of Fatty Acid and Sterol Carrier Proteins: Effects of Ethanol. *Biochemistry* **1995**, *34* (37), 11919–11927.

(6) Taylor, B. N.; Kuyatt, C. E. *Guidelines for Evaluating and Expressing the Uncertainty of NIST Measurement Results*; US Department of Commerce, Technology Administration, National Institute of …, 1994; Vol. 1297.

(7) Cer, R. Z.; Mudunuri, U.; Stephens, R.; Lebeda, F. J. IC 50-to-K i: A Web-Based Tool for Converting IC 50 to K i Values for Inhibitors of Enzyme Activity and Ligand Binding. *Nucleic Acids Res.* **2009**, *37* (suppl_2), W441–W445.

(8) García, F. L.; Szyperski, T.; Dyer, J. H.; Choinowski, T.; Seedorf, U.; Hauser, H.; Wüthrich, K. NMR Structure of the Sterol Carrier Protein-2: Implications for the Biological Role. *J. Mol. Biol.* **2000**, *295* (3), 595–603.

(9) Kay, L. E.; Ikura, M.; Tschudin, R.; Bax, A. Three-Dimensional Triple-Resonance NMR Spectroscopy of Isotopically Enriched Proteins. *J. Magn. Reson.* **1990**, *213* (2), 423–441. https://doi.org/10.1016/j.jmr.2011.09.004.

(10) Wittekind, M.; Mueller, L. HNCACB, a Highsensitivity 3D NMR Experiment to Correlate Amideproton and Nitrogen Resonances with the α-Carbon and β-Carbon Resonances in Proteins. *J. Magn. Reson. Ser. B* **1993**, *101*, 214–217.

(11) Grzesiek, S.; Bax, A. Correlating Backbone Amide and Side Chain Resonances in Larger Proteins by Multiple Relayed Triple Resonance NMR. *J. Am. Chem. Soc.* **1992**, *114* (16), 6291–6293.

(12) Delaglio, F.; Grzesiek, S.; Vuister, G. W.; Zhu, G.; Pfeifer, J.; Bax, A. NMRPipe: A Multidimensional Spectral Processing System Based on UNIX Pipes. *J. Biomol. NMR* **1995**, *6* (3), 277–293. https://doi.org/https://doi.org/10.1007/BF00197809.

(13) Lee, W.; Tonelli, M.; Markley, J. L. NMRFAM-SPARKY: Enhanced Software for Biomolecular NMR Spectroscopy. *Bioinformatics* **2015**, *31* (8), 1325–1327. https://doi.org/10.1093/bioinformatics/btu830.

(14) Passaro, S.; Corso, G.; Wohlwend, J.; Reveiz, M.; Thaler, S.; Somnath, V. R.; Getz, N.; Portnoi, T.; Roy, J.; Stark, H. Boltz-2: Towards Accurate and Efficient Binding Affinity Prediction. *BioRxiv* **2025**.

(15) Mirdita, M.; Steinegger, M.; Söding, J. MMseqs2 Desktop and Local Web Server App for Fast, Interactive Sequence Searches. *Bioinformatics* **2019**, *35* (16), 2856–2858.

(16) Jackson, T. W.; Scheibly, C. M.; Polera, M. E.; Belcher, S. M. Rapid Characterization of Human Serum Albumin Binding for Per-and Polyfluoroalkyl Substances Using Differential Scanning Fluorimetry. *Environ. Sci. Technol.* **2021**, *55* (18), 12291–12301.

(17) Zhang, L.; Ren, X.-M.; Guo, L.-H. Structure-Based Investigation on the Interaction of Perfluorinated Compounds with Human Liver Fatty Acid Binding Protein. *Environ. Sci. Technol.* **2013**, *47* (19), 11293–11301.

(18) Ren, X.-M.; Qin, W.-P.; Cao, L.-Y.; Zhang, J.; Yang, Y.; Wan, B.; Guo, L.-H. Binding Interactions of Perfluoroalkyl Substances with Thyroid Hormone Transport Proteins and Potential Toxicological Implications. *Toxicology* **2016**, *366*, 32–42.

(19) Weiss, J. M.; Andersson, P. L.; Lamoree, M. H.; Leonards, P. E. G.; Van Leeuwen, S. P. J.; Hamers, T. Competitive Binding of Poly-and Perfluorinated Compounds to the Thyroid Hormone Transport Protein Transthyretin. *Toxicological sciences* **2009**, *109* (2), 206–216.

(20) Khazaee, M.; Christie, E.; Cheng, W.; Michalsen, M.; Field, J.; Ng, C. Perfluoroalkyl Acid Binding with Peroxisome Proliferator-Activated Receptors α, γ, and δ, and Fatty Acid Binding Proteins by Equilibrium Dialysis with a Comparison of Methods. *Toxics* **2021**, *9* (3), 45.

(21) Shen, Y.; Bovee, T. F. H.; Molenaar, D.; Weide, Y.; Nolles, A.; Braucic Mitrovic, C.; van Leeuwen, S. P. J.; Louisse, J.; Hamers, T. Optimized Methods for Measuring Competitive Binding of Chemical Substances to Thyroid Hormone Distributor Proteins Transthyretin and Thyroxine Binding Globulin. *Arch. Toxicol.* **2024**, *98* (11), 3797–3809.

(22) Yang, D.; Han, J.; Hall, D. R.; Sun, J.; Fu, J.; Kutarna, S.; Houck, K. A.; LaLone, C. A.; Doering, J. A.; Ng, C. A.; Peng, H. Nontarget Screening of Per-and Polyfluoroalkyl Substances Binding to Human Liver Fatty Acid Binding Protein. *Environ. Sci. Technol.* **2020**, *54* (9), 5676–5686.
